# Fertility decline in *Aedes* mosquitoes is associated with reduced maternal transcript deposition and does not depend on female age

**DOI:** 10.1101/2023.12.05.570316

**Authors:** Olayinka G. David, Andrea V. Arce, Andre Luis Costa-da-Silva, Anthony J. Bellantuono, Matthew DeGennaro

## Abstract

Female mosquitoes undergo multiple rounds of reproduction known as gonotrophic cycles. A gonotrophic cycle spans the period from bloodmeal intake to egg laying. Nutrients from vertebrate host blood are necessary for completing egg development. During oogenesis, a female pre-packages mRNA into her oocytes, and these maternal transcripts drive the first two hours of embryonic development prior to zygotic genome activation. In this study, we profiled transcriptional changes in 1-2 hours old *Aedes aegypti* embryos across two gonotrophic cycles. We found that homeotic genes which are regulators of embryogenesis are downregulated in embryos from the second gonotrophic cycle. Interestingly, embryos produced by *Ae. aegypti* females progressively reduced their ability to hatch as the number of gonotrophic cycles increased. We show that this fertility decline is due to increased reproductive output and not the mosquitoes’ age. Moreover, we found a similar decline in fertility and fecundity across three gonotrophic cycles in *Ae. albopictus*. Our results are useful for predicting mosquito population dynamics to inform vector control efforts.

## INTRODUCTION

Anautogenous mosquitoes rely on nutrients from vertebrate host blood to complete oogenesis [1]. Anthropophilic species such as *Aedes aegypti* and *Anopheles gambiae* are competent vectors of deadly pathogens causing over 600,000 human deaths annually [2]. Under optimal conditions, female *Ae. aegypti* mosquitoes can undergo multiple reproductive cycles also known as gonotrophic cycles [3]. To fulfil each gonotrophic cycle, female mosquitoes require a separate bloodmeal intake. Transmission of pathogens by mosquitoes typically requires successive blood feeding events or gonotrophic cycles [4], although some vertical transmissions have been reported [5]. Vertebrate host blood is a critical activator of vitellogenesis, the maturation stage of egg development associated with high metabolism and uptake of egg yolk components [6,7], and maternal transcript synthesis [8]. These maternally derived transcripts are maintained in the mature oocyte until they are required to drive developmental processes during early embryogenesis [9,10].

Homeotic genes or homeoboxes (Hox) belong to a family of genes that encode transcription factors which are critical regulators of early embryonic development [11,12]. Hox genes encode homeodomain proteins with a characteristic DNA-binding structural profile for controlling the expression of target downstream genes during development [13–16], and so confer a positional identity and specification during embryonic segments formation [17,18]. In addition to their role in body patterning during early development, Hox genes also function in organogenesis by activating genes within target cell lineages required for organ formation during embryogenesis [19,20]. Many homeotic loci have been identified and extensively studied in *Drosophila* development [11,12,21,22]. A classic example is the *bicoid* (*bcd*) protein which serves as a positive transcriptional regulator of the segmentation gene, *hunchback* (*hb*), to determine presumptive poles in the early embryo in a concentration dependent manner [23,24]. Although several *Drosophila* homeodomain proteins are maternally encoded, they act to differentially activate zygotic gene expression in early development [25–29]. Studies in the malaria vector, *An. gambiae*, have identified a major difference in the genomic organization of mosquito and *Drosophila* hox gene clusters [30,31]. Whereas the *An. gambiae hox* genes are organized in a single cluster, *Drosophila* hox gene cluster is partitioned into two [30–32]. Nevertheless, multiple line of evidence suggests a conserved role for *hox* genes due to their shared DNA sequence and protein structure similarities from fly to human [30,31,33–35]. Although mosquito orthologs of some *Drosophila* maternal effect genes have been identified and used to drive germline specific expression of transgenes in mosquito embryos [36,37], more research is needed to better inform our understanding of the specific function of *hox* genes in mosquito development.

In this study, we compared the transcriptional profile in early *Ae. aegypti* mosquito embryos across two gonotrophic cycles. Using differential gene expression analysis, we identified key homeotic genes that are downregulated in the second gonotrophic cycle embryos. Interestingly, we found a continuous decline in fecundity, the total number of eggs laid, and fertility, the fraction of hatched eggs, across three gonotrophic cycles of *Ae. aegypti* and *Ae. albopictus* mosquitoes. These results suggest that maternal transcript levels of key developmental genes influence female mosquito fertility by regulating development to promote successful hatching of larvae.

## METHODS

### Mosquito rearing

All mosquitoes used in this study were reared in the insect facility at 27 ± 1°C and 70 ± 10% relative humidity in a 14:10 light-dark cycle with lights on at 8 a.m. For colony maintenance, eggs were hatched in a 0.5 L vacuum sealed Mayson jar containing preboiled deoxygenated deionized water with dissolved half tablet TetraMin tropical fish food. Twelve to twenty-four hours post hatching, ∼250 larvae were transferred into a rearing pan containing 2 L of deionized water (DI) and fed Tetramin tablets until they pupated. Pupae were transferred to a porcelain ramekin containing DI water and allowed to emerge in an insect rearing cage. Adult mosquitoes were maintained in a 1:1 male to female ratio and provided 10% sucrose solution *ad libitum*.

Upon emerging, adult mosquitoes were allowed to freely mate for 3-5 days. Females were then provided with defibrinated whole sheep blood supplemented with 0.4 mM ATP using a glass feeder (# 1588-50, NDS Technologies, Vineland, NJ) covered in a thin parafilm that has been pre-rubbed on human arm to enhance female attraction and connected to a water bath maintained at 37°C. Three to four days post blood feeding, Whatman laden ramekin with DI water was placed in the rearing cage to serve as oviposition substrate for egg laying. Following egg laying, egg papers were kept damp for additional 5-7 days to allow full embryonation before being transferred to a storage bag.

### Fecundity and fertility assays

To determine fecundity, the number of eggs deposited per female, 5–7-day old, mated females were starved on water overnight and then provided with a blood meal to repletion as described above. Fifty engorged females were transferred to a fresh cage and maintained on 10% sucrose solution. 72-96hrs post blood meal, females were cold-anaesthetized and individually prepared for oviposition in a 24-well plate as previously described [38]. Females were allowed to lay eggs over a period of 24hrs and individuals that failed to lay eggs were removed from the experiment. Upon egg collection, females were returned into the rearing cage with unlimited access to 10% sucrose solution and allowed to recover for 24hrs after which they are prepared for the second round of egg collection. Egg papers were kept moist for additional 3-5 days post collection and the number of eggs laid by individual female was subsequently counted. To determine fertility, the percentage of eggs that hatched into larvae, 5–7-day old eggs were hatched in individual 50mL cups containing DI water with dissolved TetraMin powder over a period of 3 days after which hatched larvae were counted. The same protocol was used in all experiments except in the delayed gonotrophic assay where some females were provided with their first blood meal at 21 days old. All experiments were conducted in triplicates with different mosquito rearing cohorts.

### Transcriptomic data processing and analysis

RNA sequencing (RNA-seq) data for 1-2hrs old wild-type *Ae. aegypti* embryos was retrieved from the transcriptome reported in our previous study [3]. Each library read was trimmed to remove adapter sequence and then mapped to gene models from AaegL5.0 genome [39,40]. Upon mapping, transcript level abundance was determined from mapped reads using tximport [41]. Principal component and differential gene expression analyses were performed on read counts using R’s shiny DEBrowser [42]. Differential expression was calculated using Deseq2 at corrected FDR of α < 0.01. Gene ontology (GO) enrichment was performed on differentially expressed genes at α < 0.01 using the bioinformatics resources on vectorbase.org, and redundant or obsolete GO terms were removed.

## RESULTS

### Continuous fertility and fecundity decline over three gonotrophic cycles in Aedes mosquitoes

Our previous study showed a fecundity and fertility decline between the first and second gonotrophic cycles in *Ae. aegypti* mosquitoes [3]. To determine if this phenotype is progressive and/or specific to *Ae. aegypti*, we assayed fecundity and fertility across three gonotrophic cycles in *Ae. aegypti* and *Ae. albopictus* to assess the number of eggs females laid, and the fraction of those eggs that hatched into viable larvae, respectively (figure 1*a*). We found a continuous decrease in fertility and fecundity across three gonotrophic cycles in *Ae. aegypti* (figures 1*b &* 1*c*) and *Ae. albopictus* (figures 1*d &* 1*e*), suggesting that this effect may be conserved across *Aedes* species.

**Figure 1.**
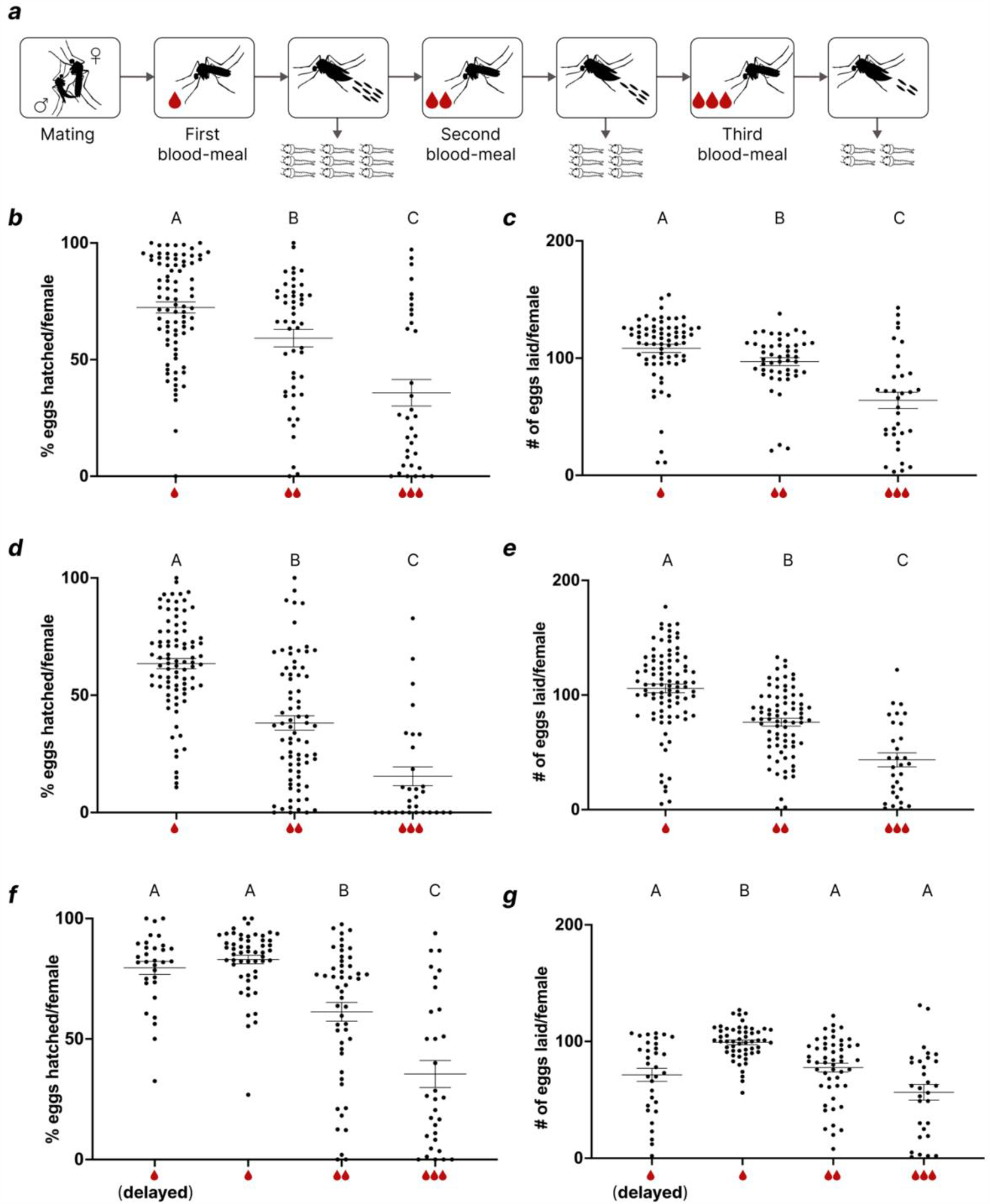
Fertility and fecundity decline in *Aedes* mosquitoes following the second and third gonotrophic cycle. (*a*) Graphical illustration of mosquito fecundity and fertility assays over three gonotrophic cycles (*b*) Percentage of individual female *Ae. aegypti* eggs hatched over three gonotrophic cycles (*n*=34-65, *p*<0.0001). (*c*) Number of eggs laid by individual *Ae. aegypti* female over three gonotrophic cycles (*n*=34-70, *p*<0.0001). (*d*) Percentage of eggs hatched per individual *Ae. albopictus* female over three gonotrophic cycles (*n*=30-87, *p*<0.0001). (*e*) Number of eggs laid by individual *Ae. albopictus* female over three gonotrophic cycles (*n*=30-91, *p*<0.0001). (*f*) Percentage of eggs hatched per individual *Ae. aegypti* female over three gonotrophic cycles in delayed gonotrophic cycle assay (*n*=30-54, *p*<0.0001). (*g*) Number of eggs laid by individual *Ae. aegypti* female over three gonotrophic cycles in delayed gonotrophic cycle assay (*n*=30-54, *p*<0.0001). Statistical analysis was performed using Kruskal-Wallis test followed by Dunn’s multiple comparison test. For all dot plots, each dot represents data from an individual female, long lines represent the mean, and short lines represent standard error. Each experiment was performed using three mosquito cohorts of 24 females. Groups marked with different letters are significantly different.

To exclude the age of females as the determinant of fertility and/or fecundity decline, we performed a delayed gonotrophic cycle assay in which experimental females were given their first blood meal 20 days post eclosion while control females were being provided a third blood meal. We found no significant difference in fertility between the regular and delayed gonotrophic mosquito groups during the first gonotrophic cycle (figure 1*f*). However, there was a significant difference in fecundity between the two groups in the first gonotrophic cycles (figure 1*g*). These results suggest that the age of the mosquitoes does not cause the hatching reduction seen in embryos produced in the second and third gonotrophic cycles. Rather, it suggests that increased reproductive output may cause a fertility decline in females.

### Transcriptomes of early Ae. aegypti embryos revealed differential expressions between gonotrophic cycles

Pre-zygotic transition *Ae. aegypti* embryo gene expression analysis showed transcriptional changes cluster by gonotrophic cycle [3], suggesting that maternal mRNA deposited during oocyte development may vary between reproductive cycles. To gain better understanding into this hypothesis, we analyzed our transcriptomic data to identify changes in maternal gene expression profile between the first and second gonotrophic cycle *Ae. aegypti* embryos (figure 2*a*). To assess cluster patterns across samples, we performed principal component analysis (PCA) on library reads from the first and second gonotrophic cycles embryos. PCA revealed sample clusters by gonotrophic cycle (figure 2*b*). Next, we performed differential expression analysis between gonotrophic cycle embryos using Deseq2 and identified a total of 267 differentially expressed genes (DEGs) (figure 2*c*). Out of the DEGs, we found that more than three times as many were downregulated (204) in the second gonotrophic cycle embryos than were upregulated (63) (figure 2*c*).

**Figure 2.**
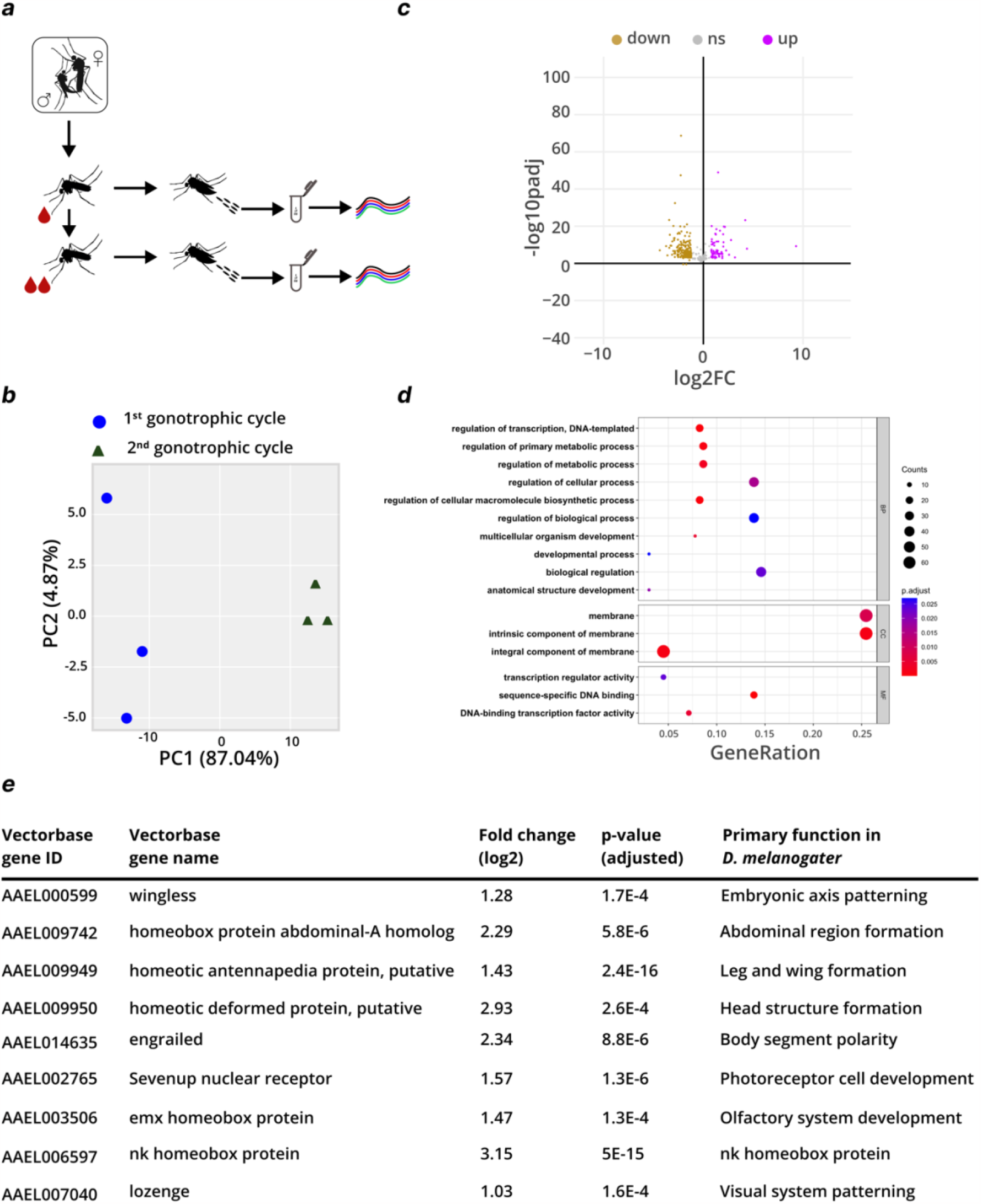
Homeobox genes are downregulated in *Ae. aegypti* pre-zygotic transition embryos produced in the second gonotrophic cycle. (*a*) Schematic of sample collection for RNA-seq analysis. (*b*) Principal component analysis of first and second gonotrophic cycle pre-zygotic *Ae. aegypti* embryo transcriptomes. (*c*) Volcano plot of differentially expressed pre-zygotic *Ae. aegypti* embryo genes between the first second gonotrophic cycles, (*n*=3, FDR<0.01). (*d*) Gene ontology enriched terms among differentially expressed genes in (*c*), (FDR<0.05). (*e*) Maternally encoded genes downregulated in the second gonotrophic cycle embryos in (*c*), and their primary function in *D. melanogaster*. See also supplemental files.

### Maternally derived hox genes are downregulated in the second gonotrophic cycle embryos of Ae. aegypti mosquitoes

We next performed a gene ontology enrichment assessment to determine the biological processes, cellular components and molecular functions significantly represented among the differentially expressed gene sets. Interestingly, regulators of cellular processes and development are highly represented across the three functional trait classes (figure 2*d*). We identified maternally encoded regulators of embryonic development, including 7 members of the highly conserved *hox* gene family, downregulated in the second gonotrophic cycle embryos (figure 2*e*). Together, these results show transcriptional changes in maternal mRNA during the early development of mosquito embryos are associated with the number gonotrophic cycles a female mosquito has experienced.

## DISCUSSION

We previously reported a maternal effect decline in *Ae. aegypti* mosquito fertility associated with changes in the transcriptome of early embryos that resulted from the mutation of an olfactory coreceptor gene, *orco* [3]. Here, we provide evidence for the role of additional maternal gene products in sustaining mosquito fertility over multiple reproductive cycles. Using a differential gene expression analysis of maternally derived genes in the first and second gonotrophic cycle embryos, we identified 204 genes that are downregulated in the second reproductive cycle *Ae. aegypti* embryos. Among downregulated genes are 9 regulators of early embryogenesis—seven of which are members of the *hox* gene family that promote morphogenesis and organogenesis during early development [11,12], and two non-homeotic genes involved in embryonic body axis formation [43] and photoreceptor cell fate determination [44]. Downregulated homeobox protein encoding genes include regulators of leg and abdominal region formation [45,46], morphological specialization of head segments [47,48], wing morphogenesis [49], and visual system development [50]. Interestingly, *hox* dosage and differential expression levels of partner *hox* genes have been associated with morphogenesis in *Drosophila* embryos [51,52]. Furthermore, the localized expression pattern characteristic of many homeotic and maternal effect genes coupled with their strict expression level requirement for function, as demonstrated by RNAi and genetic manipulation experiments in other insects, suggest that downregulating the expression of key developmental genes could have phenotypic effects during embryogenesis [24,55].

To determine and compare the viability of eggs produced during different reproductive cycles, we assessed the fertility and fecundity of *Aedes* mosquitoes across three gonotrophic cycles. Interestingly, both fertility and fecundity declined continuously across the three reproductive cycles in *Ae. aegypti* and *Ae. albopictus*. Given our previous report that *Ae. aegypti* females store sperm that could last them more than three rounds of reproduction, and that over 90% of eggs laid in the second gonotrophic cycle are fertilized [3], it is unlikely that the fertility decline was due to sperm deficiency. Rather, we propose that a reduced deposition of key maternal gene products during oogenesis could negatively impact on embryonic development and/or hatching. Moreover, results from our delayed gonotrophic cycle experiment show that the fertility decline phenotype, and perhaps the reduced maternal transcripts in the second reproductive cycle embryos, may result from increased reproductive output and is independent of females’ age. The impacts on female mosquito fertility we have shown could be explained by depletion of nutrient reserves when oogenesis occurs multiple times.

Female mosquitoes’ reproductive output has been positively correlated with nutritional reserves during the previtellogenic resting stage of oogenesis [56,57]. Fertility and fitness have been linked in insects, particularly in regards to immunity [58,59]. Artificial selection experiments in *Drosophila* showed that selecting for increased lifespan extends adult survivorship but decreased early reproduction and vise-versa [60]. For example, knockout female flies for the insulin-like receptor gene (*dInR*) are infertile due to non-vitellogenic ovaries, resulting in extended lifespan [61]. The decline in female fertility in our study could be a reproductive trade-off leading to reduced maternal investment during egg development to increase female mosquito survival over time.

